# Consumption of Ciprofloxacin as the sole Carbon and energy source and Phenomenal Transformation to a Culturable nanometer-sized-bacterium demonstrated by a hospital-waste-water isolate *Klebsiella* sp. SG01

**DOI:** 10.1101/2024.11.20.624549

**Authors:** Sriradha Ganguli, Ranadhir Chakraborty

## Abstract

Antimicrobial resistance is a global crisis. Biodegradation by bacteria is an effective strategy to remove the micropollutant from the environment. In this study, we demonstrate that a persistent fluoroquinolone, ciprofloxacin (CIP) can be degraded by a multidrug-resistant *Klebsiella* sp. SG01 and used as its only carbon source. The degradation was quantified using UV-vis spectroscopy and the degraded product was less toxic than the parent compound as tested against a susceptible *Escherichia coli* K12. SG01 changes into nano-sized cells as culturable nanobacterium, passes through a 0.22 μm pore-size filter while growing on ciprofloxacin, and shows a shorter generation time than cells grown on glucose or rich medium. The basis for the changed growth phenotype of nano-SG01 cells and metabolic changes was partially established by the whole-genome transcriptome.

**Importance:** Fluoroquinolones (FQs) constitute a class of persistent antimicrobials, having a high affinity for sludge, sediments, and soil with reported half-lives of 10.6 days in surface waters and up to 580 days in soil matrices. However, more than 50% consumption or degradation of CIP at higher concentration (∼2g/L) within 48 hours is not yet reported. Also, the isolate transforms into a nano-form to consume the antibiotic and economizes its metabolism to degrade CIP. From the One Health Approach perspective, nanobacterial transformation is an alarming concern as such viable nanobacteria continue to proliferate and may trigger infection in host. The strategy should be to utilize this powerful strain for the biodegradation of CIP.

## Introduction

One of the crucial advancements in medicine is the discovery of antibiotics. However, the rise and proliferation of antibiotic resistance (AR) have significantly compromised their effectiveness. Globally there has been an increase in antimicrobial consumption by 11.2 % (1) and is projected 200% by 2030 (2), with antibiotics covering 83% of the antimicrobials. Studies indicate that almost 70% of the antibiotics we consume are not metabolized and excreted as the parent molecule through feces. The Wastewater treatment plants (WWTPs) are inefficient in removing the antibiotic residues and they find their way into the environment and create selective pressure for the enrichment of antibiotic resistance genes (ARGs) (3). India ranks third in supplying antibiotics (20%) and 50% of vaccines to the world demand emerging as the “Pharmacy of the Global South” (4)

Fluoroquinolones (FQ) are synthetic antibiotics widely prescribed for human and veterinary healthcare. CIP is a second-generation FQ extensively prescribed against Gram-positive and Gram-negative bacterial infections (5). CIP is a recalcitrant compound that is hard to degrade due to the presence of the C-F bond and high electronegativity of fluorine along with its unique physical and chemical properties which also enhances CIP’s biological efficacy (6). In the environment, CIP was detected in agricultural soils, freshwater ecosystems, manure, and urban sludge in the range of 5μg/kg-45.5 mg/kg (7, 8). Various physico-chemical methods are employed for the removal of CIP such as photolysis, advanced oxidation, and electrochemical oxidation but these are not very efficient and cost-friendly. Also, they generate harmful metabolites through incomplete degradation leading to secondary pollution (9).

Biodegradation of CIP by several bacterial strains was conducted, such as *Labrys portucalensis* F11 which could degrade a wide range of FQs in the presence of additional carbon sources (10), *Thermus sp*. C419(11), and *Bradyrhizobium* sp.GLC_01 (12) and *Paraclostridium* sp (13). In 2024, two strains isolated from pharmaceutical wastewater, *Stutzerimonas stutze*ri, and *Exiguobacterium indicum*, were reported to degrade 35% and 28% respectively of the total CIP used in 10 days (14). In all of these degradations studies, CIP concentration ranged from 20 to 100 mg/L. In the present study, CIP at a concentration of 2 g/L as the sole carbon source has been used.

We have isolated a strain *Klebsiella* sp SG01 capable of degrading 57% of total CIP content in the medium within 48 h of incubation. In doing so, the strain undergoes a morphological transformation into a nano-form. Nanobacteria have sparked controversy and excitement amongst researchers since their discovery. While some consider them to be the tiniest forms of life on Earth (15, 16), others put a dead end to this concept by calling them sub-microscopic balls of proteins and minerals (16). Our work turned the cartwheel and brought a paradigm shift in the understanding of CIP degradation and utilization as the sole carbon and energy source by the SG01’s ‘nano-form’ population. The present work has indicated that nanobacteria may not be confined to specific phyla, bearing small genomes and large generation times (17, 18). The probable pathway operating in SG01 to utilize CIP as the sole carbon and energy source has been proposed.

## Results

### Enrichment

An acclimation strategy was adopted to allow the enrichment of strain(s) capable of degrading CIP and eliminating non-degraders. Hence, after each acclimating cycle chances of survival of organisms that could utilize CIP as a carbon source increased. At the end of the fourth cycle, 10^9^ cells were obtained. They were clonally purified and found morphologically identical colonies and their constituent cells.

### Morphological, Biochemical, and Genomic Characterization of the isolate

The isolated strain SG01 is Gram-negative and rod-shaped. The strain is found to be a catalase-positive and oxidase-negative bacterium capable to grow in mineral salts medium without supplementation of nitrogen source, and utilizing a wide range of sugars such as lactose, sucrose, mannose, inositol, glycerol, raffinose, and dextrose (Table S1, S2). The antibiotic susceptibility profile of the strain confirms it a multidrug resistant bacterium (Table S3). Sequencing and *de novo* assembly results showed that the genome length of SG01 was 5,583,874 bp and G+C content of 57 %. The whole genome encodes a total of 5229 CDS, 77 tRNA, 3rRNA, and 1 tmRNA genes. In addition, the whole genome contains 242 phage genes. This whole genome was successfully submitted to NCBI under accession no JAQSIQ000000000. Evolutionarily the strain forms a branch from *Klebsiella pneumonniae* subsp. *pneumonia* DSM 30104 AJJID00000000 (Fig S1).

### Growth characteristics of SG01 in MSM-CIP medium in comparison to its growth in MSM-glucose and LB medium

Growth of SG01 in mineral salt medium (MSM) containing CIP (2g/L) as the sole source of carbon and energy was monitored in terms of an increase in viable cell count. In MSM-CIP, the SG01cells grew exponentially reaching 10,000 times its original cell count at 20 h of incubation. For comparison, prototrophic and heterotrophic growth of SG01 was measured in MSM supplemented with glucose (5 g/L), and Luria Bertani (LB) medium respectively. In LB, cells continued to grow exponentially with a 1000 times increase in cell number at 8 h of incubation; on the other hand, the stationary phase was attained after 9 h after a 100-times increase in CFU/ml when cultured in MSM-glucose (5g/L) (Fig.1. a-c).

### Quantification of CIP in spent MSM-CIP medium using UV-vis Spectroscopy

Since its functional groups have been protonated, pure ciprofloxacin diluted in MSM medium at pH 4.0 (pH adjusted with 0.1 N HCl) is largely in a cationic form, which has broadened the absorption spectra and produced three absorption maxima at 315 nm, 326 nm, and 332 nm, respectively. Linearity for the spectrophotometric method at 315 nm was noted over a concentration range of 2-20 μg/ mL with a correlation coefficient of 0.992 (Fig S2).

A standard curve was plotted depending on the best-fit curve (R2 value=0.992) at 315nm. We observed a 57 % reduction in CIP concentration after 54 h of incubation. Interestingly, during the lag phase cells initiated degradation, approximately 27% reduction in CIP concentration was observed after 6 h of incubation (Fig. 1d).

**Figure 1.**
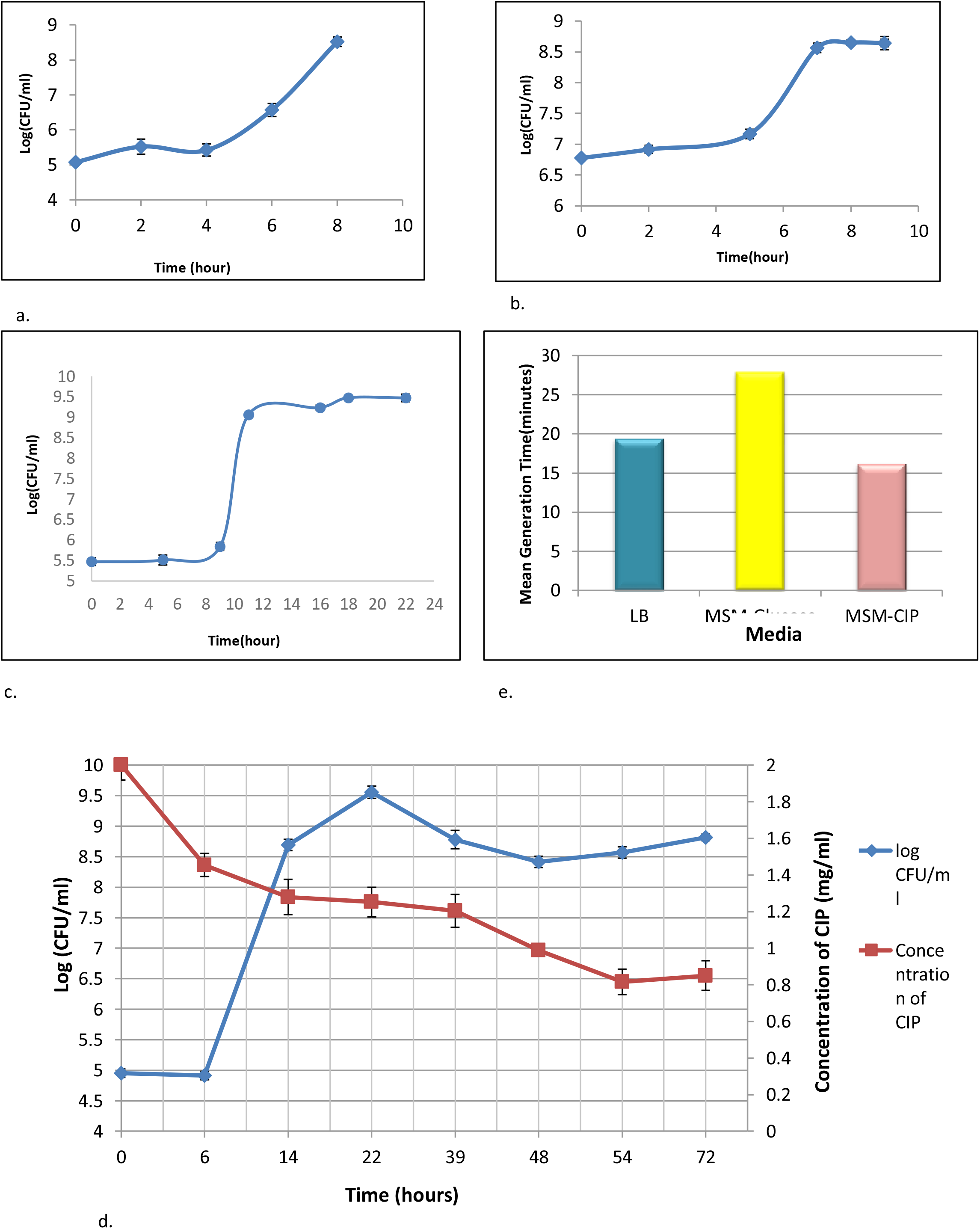
Growth studies of SG01 in different media. CFU/mL was measured at different time interval. Results are outcome of three biological replicates. a, Growth assay in Luria Bertani (LB); b, Growth assay in MSM-Glucose (5g/L); c, Growth assay in MSM-CIP (2g/L); d, Growth assay in MSM-CIP and quantification of CIP using UV-vis Spectrophotometer; e, Mean generation time of SG01 in different media.

### Residual antibacterial activity of the degraded CIP in the culture medium

The residual antibacterial activity of the degraded CIP was checked against an antibiotic-susceptible wild-type bacterium, *Escherichia coli* K12. The zone diameter reduced with fixed volume of culture filtrate collected from aging culture in terms of incubation hours, suggesting that the degraded CIP due to bacterial activity is less toxic as compared to the parent molecule (Fig S3).

### Cellular plasticity

During growth assays, it was observed that turbidometric measurement, which is used to indirectly determine an increase in cell number, did not apply to SG01-cells grown in CIP; even in cases where they were present in significant numbers (determined by standard plate count), optical density (OD) measurements could not be used to detect growth. Log phase cultures of SG01 grown in various media were aseptically passed through a bacterial filter to demonstrate the phenomenon of size reduction and that refractoriness to OD measurements may only occur if the cells are below the size threshold that significantly scatters light. The number of viable bacteria present in the unfiltered culture and filtrate was measured using a standard plate count method. Only ≈ 0.01% of cells grown in MSM-glucose or LB were able to pass through the bacterial membrane filter (pore size, 0.22 μm), compared to over 98% of cells produced in MSM-CIP. Therefore, a ‘t’ test comparing the means of the two groups, the viable cell count obtained from the unfiltered and filtrate, revealed that a significantly smaller number of cells produced in MSM-glucose (P = 0.0207) or LB (P = 0.025) could pass through the bacterial filter. Conversely, there was no statistically significant variation in the number of cells found in the unfiltered and filtrate of cultured cells in MSM-CIP (P = 0.914). This suggests that the cells produced in CIP will be less than 0.22 μm in size. Negative staining with Nigrosin produced a distinct image of nanoscale cells (Fig.2). Additionally, the average length of each cell growing in MSM-glucose and MSM-CIP was found to be 1.35 μm and 0.16 μm (160 nm), respectively, according to scanning electron microscopy (Fig.S4 a,b).

**Figure 2.**
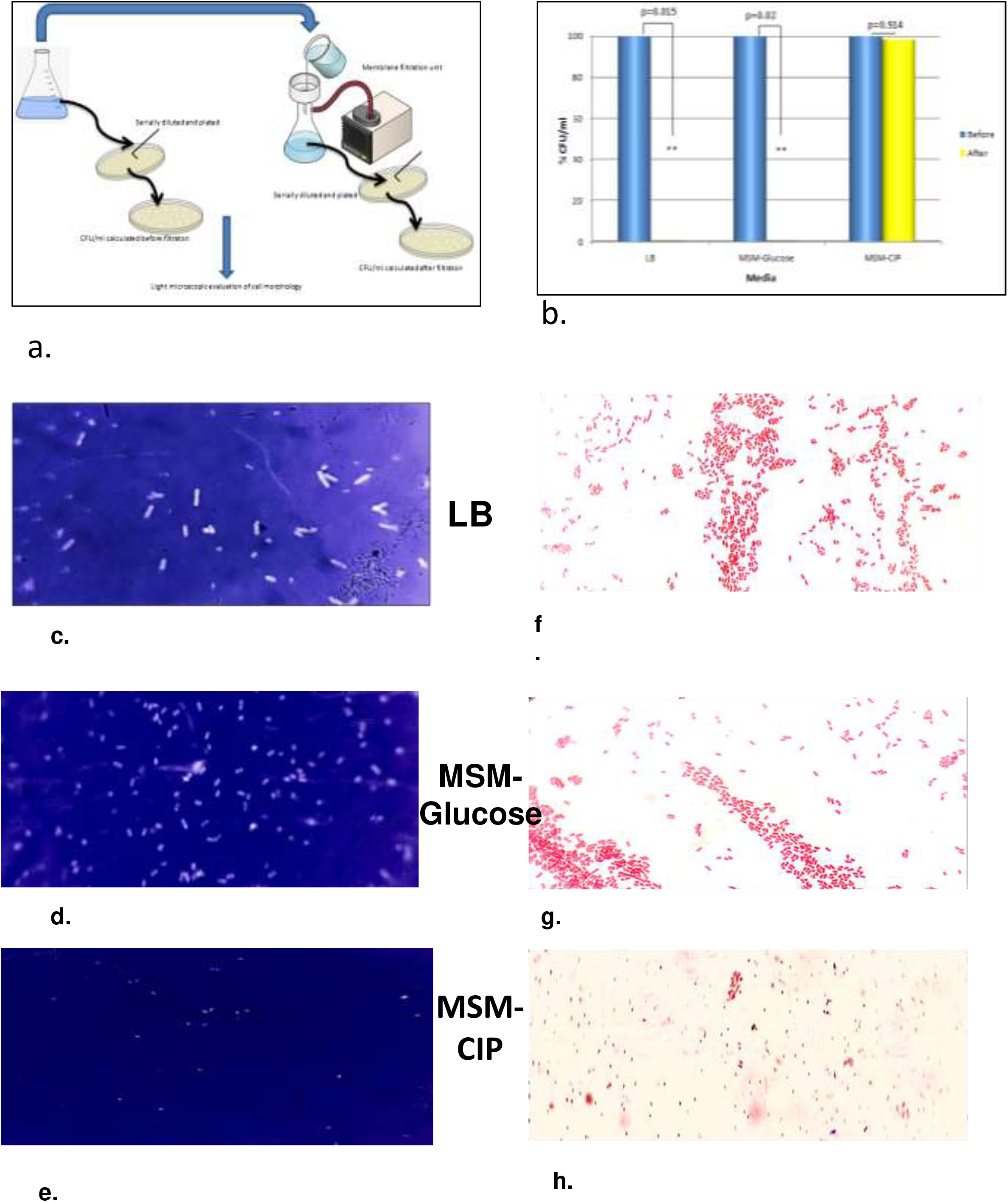
Revelation of reduction in cell dimensions from micrometer to nanometer-sized bacterium. a, Schematic illustration of passing SG01 culture grown cells in different media through the bacterial membrane filtration unit followed by ascertaining the cell counts by standard plating technique; b, Quantification of cells (%) grown in different media, before and after passing through the membrane filter; **cells grown in LB and MSM-glucose showed a significant difference between filtered and unfiltered cells; cells grown in MSM-CIP showed no significant difference in the filtered and unfiltered sample; c, d, e, Negative staining images (light microscopic) of cells grown in LB, MSM-glucose, MSM-CIP; f,g, h, Images (light microscopic) of Gram staining of cells grown in LB, MSM-glucose, MSM-CIP.

### Transcriptome analysis

The RNA-seq profiling of *Klebsiella* sp SG01 while growing with CIP as the sole carbon source, and without CIP but supplemented with glucose as the sole carbon source was performed to unveil the genes intricate in the degradation process. A total of 5367 differentially expressed genes (DEGs) were generated via mapping of reads to the reference genome. The comprehensive information on genes and their annotations are given in Supplementary Excel File 1. Based on the level of significance (p<0.05), 1637 genes were upregulated and 2232 genes were downregulated. Among the upregulated genes, 163 genes were significantly upregulated and 259 genes were significantly downregulated (log 2FC >2). Hence, more genes were significantly downregulated as compared to upregulation (Supplementary Excel file 1).

The expression of genes coding for peroxidase-related enzymes in CIP-grown cells was significantly high (log 2FC=2.72) compared to glucose-grown cells. The superoxide-generating agents also triggered the expression of *soxR* and *soxS* which are the activators of superoxide stress response genes (Li et al., 1994). However, there was a downregulation of superoxide dismutase genes *sodC* (−2.56 log 2FC) and *sodB* (−1.97 log2FC). DEGs related to the transporters were analyzed; approximately 64 transporters (log 2FC >1) were upregulated in CIP-fed cells. Fifteen transporter genes expressed with log2 fold changes of ≥2 to 5 in CIP-grown cells. Log2 fold change was highest (5.044) in the case of multidrug resistance (MDR) family major facilitator superfamily (MFS) transporter gene (PRL05_08210) followed by 4.79 log2 (FC) expression of glycerol-3-phosphate transporter gene (*glpT*), 4.65 log2 (FC) expression of metal ATP-binding cassette (ABC) transporter permease gene (PRL05_00415), and 4.12 log2 (FC) expression of two genes coding for metal ABC transporter substrate-binding protein (PRL05_00400) and manganese/iron ABC transporter ATP-binding protein (PRL05_00405). A log2 (FC) threshold of ≥1 to 2 was observed for 51 transporter genes which reflects meaningful changes in gene expression of CIP-grown SG01 cells. Genes responsible for fatty acid biosynthesis were mostly downregulated, the expression of *fabI,fabF, accB*, and PRL05_04495 (ketoacyl-ACP synthase III) were significantly downregulated (log 2FC <1, p<0.05). The genes for fatty acid degradation like *fadE, fadJ*, acetyl-coA-acetyltransferase, *fadL*, and *fadA*, were significantly upregulated (log 2FC>1). The *glp* operon is upregulated in CIP-grown cells.

### Network analysis and Identification of the hub genes

We computationally derived the Protein-Protein Interaction (PPI) network of the upregulated and downregulated genes. As the figure S5 (a-b) depicts, through the STRING database we could highlight the sub-clusters and their interactions. For the upregulated network, we highlighted the phosphotransferase system (PTS), fatty acid degradation, translation, stress response, and glycerol metabolism. For the downregulated, there were numerous clusters with genes for pyruvate metabolism, energy metabolism, cell morphogenesis, DNA repair, and transporters.

After analyzing the functional categories, we tried to pinpoint the “hub genes” that might play an important role in this metabolic shift. Through the CytoHubba plugin, we highlighted 10 hub genes *rpmA, rplS,rpmG, rpsQ, rplP, rpsI,rpmB, rplT, and rpsT* governing the upregulated pathways. While the *nuo* operon genes *nuoB, nuoH, nuoI nuoA, nuoF, nuoE, nuoM, nuoN, nuoJ, and nuoL* were identified as the hub genes for the downregulated pathways. Two genes *glpK* and *glpC* are identified as the two highest-scored bottleneck genes in the upregulated pathways and *nuo*CD and *eno* are the two top-scored bottleneck genes in the downregulation network. Bottleneck genes serve as a crucial link between the clusters in a network or simply they are the connecting links between networks. Hence, they do participate in regulating the metabolic pathways.

## Discussion

CIP is a broad-spectrum fluoroquinolone antibiotic, widely prescribed across the globe. Being highly persistent it poses a serious threat to the ecosystem, encouraging the proliferation of antibiotic resistance. Hence its removal from the water bodies is a serious challenge. Our study shows promising results to remove this pollutant.

The growth studies demonstrated that CIP could be the strain SG01’s only source of carbon and energy, resulting in a 1000-fold increase in cell number (Fig.1.c). Generally speaking, environmental stress would raise the concentration of reactive oxygen species, most likely leading to cell death (19). Since hydroxyl radicals are extremely reactive species that may break down a variety of chemical substances, including ciprofloxacin, it is reasonable to assume that the phenomena will benefit the surviving cells. DEGs linked to reactive oxygen species (ROS) so partially mirrored a similar situation. Previous reports have shown the role of OH radicals playing a significant role in the degradation of CIP through Fenton reactions and radiolytic decomposition (20). The downregulation of *sod* genes might be a strategy or adaptation by the strain to enhance the production of OH radicals for facilitating CIP degradation. To measure the degradation of CIP, the lag phase of SG01’s development, which lasted for six hours, was investigated (Fig. 1.d). The bacteria in the culture broke down 27% of the CIP content during the lag phase. This phase is essentially one of adaptation, during which cells were likely collecting metabolites such as fatty acids biosynthesised from carboxylic acid released by ciprofloxacin breakdown as they adjusted to the new nutritional environment (CIP as the only available carbon and energy source). In the metabolism and degradation of ciprofloxacin, the quinolone ring can undergo several transformations, including decarboxylation and other modifications, resulting in various derivatives. The piperazine moiety might also be hydroxylated by hydrolases, dioxygenases (21) supported by significant upregulation of Rieske (Fe-S) containing enzyme (PRL05_14675), and ring-opening enzymes *pobA, ygiD*. These modifications alter the structure and bioactivity of the molecule. Removal of the carboxylic acid group (−COOH) at the 3-position of the quinolone ring leads to the formation of a decarboxylated ciprofloxacin. This alteration generally results in a loss of antibacterial activity, as the carboxyl group is essential for binding to bacterial enzymes like DNA gyrase and topoisomerase IV. The possibility of generating decarboxylated-CIP in the culture medium is corroborated by the observation that CIP-susceptible wild type *E. coli* K12 has shown progressive reduction in inhibition against culture filtrate containing degraded CIP (Fig. S3). Hence, it is hypothesised that chains of fatty acids produced in the lag phase will undergo beta-oxidation to produce acetyl-Co-A in the log phase. This can then feed into pathways that make three-carbon intermediates, such as dihydroxy acetone phosphate (DHAP), which ultimately form glycerol. The genome-wide expression profile can be used to establish the sequence of events that occur during the lag phase. Hence, bacterial metabolism is rewired to extract the carbon chain from CIP, metabolizing it through the *glp* operon (glycerol metabolism). During the log phase, long-chain fatty acids are metabolized through the *fad* pathway (upregulated), generating glycerol utilization pathway intermediates (thus upregulation of the glp operon) (Supplementary Excel File 2). The glycerol being phosphorylated to glycerol-3-phosphate (*glpK*) further oxidized to DHAP by *glpB*/*glpD*.

Detecting visible growth of the strain in MSM supplemented with CIP in terms of increase in optical density was so imprecise that calculating CFU/ml over time to demonstrate the growth of nanobacterium was nearly serendipitous. This study offers up a new line of inquiry into the probable occurrence of nanobacteria in many ecological niches, as well as the impacts of these microbes that are currently unknown. The smaller size of SG01 when grown in CIP resulting in a greater surface area to volume ratio (calculations presented in the Supplementary file 1) may enable the growing cells to meet their metabolic demands through efficient uptake of nutrients. We performed a comparative transcriptome of cells grown in MSM-glucose and MSM-CIP to corroborate growth-phenotype. An expression quantifying log2 (Fold Change) of 2 or higher was chosen to focus on genes that exhibit more substantial changes. Fifteen transporter genes expressed with log2 fold changes of ≥2 to 5 in CIP-grown cells. Log2 fold change was highest (5.044)in the case of multidrug resistance (MDR) family major facilitator superfamily (MFS) transporter gene (PRL05_08210) followed by 4.79 log2 (FC) expression of glycerol-3-phosphate transporter gene (*glpT*), 4.65 log2 (FC) expression of metal ATP-binding cassette (ABC) transporter permease gene (PRL05_00415), and 4.12 log2 (FC) expression of two genes coding for metal ABC transporter substrate-binding protein (PRL05_00400) and manganese/iron ABC transporter ATP-binding protein (PRL05_00405). A log2 (FC) threshold of ≥1 to 2 was observed for 51 transporter genes which reflects meaningful changes in gene expression of CIP-grown SG01 cells (Supplementary Excel file 2).The comparative transcriptome of cells grown in MSM-glucose and MSM-CIP corroborated with the growth phenotype. The shift in cell morphology from rod-shape to spherical as seen morphologically was due to the downregulation of two rod-shape determining genes *mreB* (log 2FC= −1.15), *mreD* (log2FC = −1.01), cell division site-selection gene *minD* (log 2FC= −1.47) and *zapD* (lig 2FC=−1.5). An overexpression of (log 2FC=1.23) compensates for the effect of *zapD* and allows cell partitioning even in low concentrations of *ftsZ* (log 2FC= −0.5) (26−28). As there was size reduction, the requirement of building blocks for generating daughter cells in MSM-CIP was less, expression of genes responsible for cell wall synthesis (*daccC, mrdA, bacA*), septum site-determining gene *minD*, and cell division *zapD* were significantly downregulated along with genes for energy metabolism (*zwf, sucC, fumA,nifA, mqo*) (22). Since MSM-CIP grown SG01 cells have shown no retardation in growth rather there was a shortening in division time as expected, and expression of genes responsible for cell division remained unchanged.

Previous studies have shown that *Pseudomonas aeruginosa* treated with penicillins and carbapenems transforms from rod-shaped to spherical viable cells (23). Furthermore, the effect of antibiotics on variations in the surface area to volume (SA/V) ratio has been well studied (24). Remarkably, in contrast to traditional carbon sources like glucose or rich medium (LB), SG01 has shown faster growth with shorter generation times (g) in unorthodox carbon sources like CIP (Fig 1.e). The least ‘g’ was observed when grown CIP (16.20 min) compared to ‘g’ of 19.44 min, and 27.88 min when grown in LB and MSM-glucose respectively.

Once the environment shifts to an altered-nutrient-state, bacteria takes some time to switch on alternative metabolic pathways, causing derepression of certain operons to quickly express genes for the metabolism of alternative carbon sources (in this case, CIP), ensuring efficient adaptation to the new condition. Nevertheless, the bacterial world has evolved to convert stress to adaptation via modulating gene expressions. It is evident that under nutritional scarcity, the superfluous expenditure of energy must be prevented by regulating translation. The prime function of ribosomes is to decode the genetic machinery for protein synthesis. Apart from this, they are also the players in a large number of regulatory circuits. The translation apparatus plays a central role in cellular stresses, be it nutritional, metal, or cold. Hence, ribosomes have been termed as the switchboard for bacterial stress response (25). Previous studies demonstrated that *E*.*coli* could adapt to stresses related to cell envelope integrity by downregulating the *nuo* and *cyo* operon (26). Energy-converting NADH: ubiquinone oxidoreductase, respiratory complex I, plays an important role in cellular energy metabolism. Downregulation of *nuo* genes in bacteria would result in reduced expression of the NADH dehydrogenase complex (complex I), which is a major entry point for electrons into the respiratory chain. This downregulation can lead to a range of metabolic and physiological effects (27). With less capacity for NADH oxidation by complex I, bacteria may adapt by re-routing NADH through alternative pathways. They might upregulate other dehydrogenases that can enter electrons into the respiratory chain at different points (e.g., NDH-2, which lacks proton-pumping ability) or switch to fermentation pathways that regenerate NAD+ without using the respiratory chain. However, in anaerobic or microaerobic conditions, where alternative respiratory or fermentative pathways are more active, reduced reliance on complex I could be beneficial by minimizing energy costs associated with producing a less-needed protein complex. Since Complex I oxidizes NADH to NAD+, downregulation of *nuoCD* (one of the two top-scored bottleneck genes identified in gene networking) could disrupt the NAD+/NADH ratio. An accumulation of NADH and reduction in NAD+ levels can cause redox imbalance, affecting other metabolic pathways dependent on these cofactors, like the TCA cycle and glycolysis. Downregulating *eno* (the second top-scored bottleneck gene) presumably has influenced impaired electron transfer; more electrons may leak from the ETC, potentially interacting with oxygen to form ROS. Elevated ROS levels can help degrade ciprofloxacin. The *eno* gene encodes for the enzyme enolase, which plays a critical role in glycolysis by catalyzing genes) in bacteria and can have significant impacts on cellular metabolism and other processes, as enolase is also involved in non-glycolytic functions. Enolase’s role in glycolysis means that its downregulation would affect the balance between glycolysis and other metabolic pathways, such as the pentose phosphate pathway and TCA cycle.

Upregulation of *glpK* and *glpC* genes (highest-scored bottleneck genes in upregulated pathways) in SG01 can enhance its capacity to metabolize glycerol. *glpK* encodes glycerol kinase, the enzyme that phosphorylates glycerol to glycerol-3-phosphate. Upregulation of *glpK* increases the rate of glycerol phosphorylation, enhancing SG01’s capacity to use glycerol as a carbon and energy source. Upregulating *glpC* can enhance the flux through glycerol-3-phosphate dehydrogenase (G3PDH), which could boost the conversion of G3P to DHAP, allowing it to flow into central metabolic pathways more efficiently. Since G3PDH couples this conversion with electron transport, upregulation of *glpC* could also increase electron flux through the electron transport chain, potentially enhancing ATP production under aerobic conditions. Hence, upregulated *glpK* and *glpC* could result in more rapid growth. The increased electron flux due to *glpC* upregulation could affect cellular redox balance, potentially necessitating adjustments in other pathways for balance.

Hence, the downregulation of *nuoCD* and *eno*, and upregulation of *glpK* and *glpC*, identified as bottleneck genes in downregulated and upregulated pathways respectively (Fig. S5 a & b), satisfies the conditions to be termed as bottleneck genes that can be part of a growth strategy in MSM supplemented with ciprofloxacin as sole carbon and energy source.

The expression of these respiratory complexes was also proven to be toxic during envelope stress. Interestingly the *cpxP* is upregulated while *cpxA/R* is downregulated in CIP-fed cells. We assume the stress response to be activated during the lag phase, where *cpxA* and *cpxR* were expressed and participated in the stress-regulatory network. When stress conditions were alleviated, *cpxP* was expressed to downregulate the *cpx* response maintaining homeostasis (28).

To the best of our knowledge, we haven’t read reports of any bacteria consuming such high concentrations of CIP. This study provides a greater understanding of the ecological role of *Klebsiella* sp in the environment. The overlapping and interconnected metabolic network reveals a streamlined regulatory mechanism adopted by the strain SG01 to degrade CIP. The proposed pathway constructed (Fig.3) on the basis of growth, genome and transcriptome data, would be requiring further validation via other omics-tools to decipher the complete system biology of CIP-degradation and utilization.

**Fig. 3.**
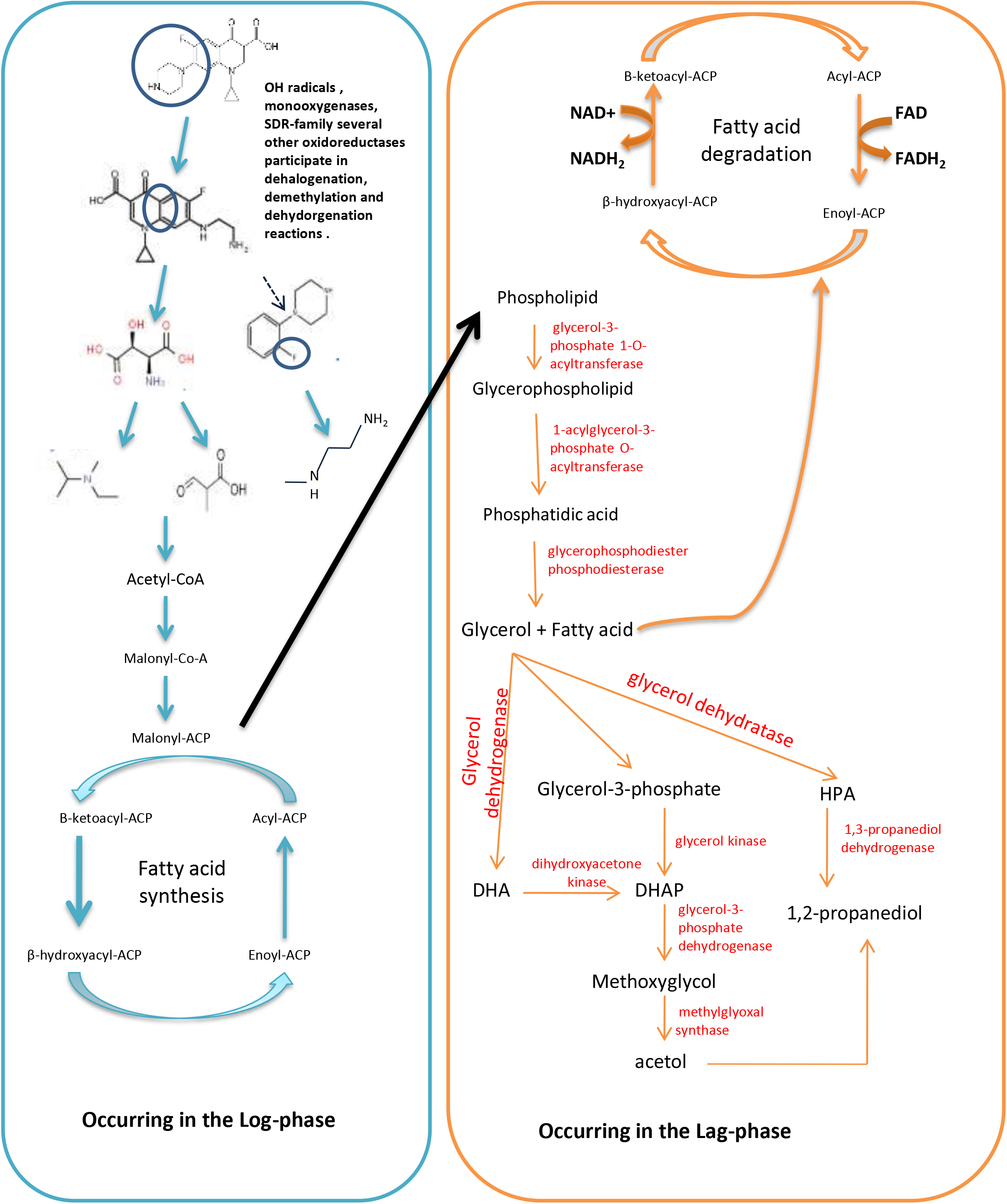
Hypothesised pathway of CIP-consumption by SG01. The left (blue) box reactions are predicted to occur during the lag phase, which feeds intermediates to the log phase metabolism (orange box on the right). CIP is initially targeted by OH radicals, oxidoreductases, hydrolases to generate acetyl-CoA which conducts the downstream metabolic pathways.

## Methods

### Enrichment culture

A waste-water sludge sample was collected from North Bengal Medical College and Hospital (NBMCH), West Bengal, India. 5 g of the sludge sample was inoculated into a 500 ml Erlenmeyer flask containing 100 ml modified mineral salts medium (MSM) supplemented with ciprofloxacin (CIP) [constituents (g/L): KH_2_PO_4_, 3.0; NaCl, 5.0; MgSO_4_. 7 H_2_O, 0.01; NH_4_Cl, 0.5; and CIP, 1.0; pH, 5.0 (adjusted with 0.1 N HCl)] and left in static condition for 7 d in the dark at 30 ^0^C. The culture (1ml) was further transferred to a fresh 100 mL MSM, supplemented with 2 g/L of CIP, and left in static condition for 7 d in the dark, at 30 ^0^C, and sub-cultured thrice under similar conditions.

### Isolation and identification of the strain capable of degrading CIP

After final sub-culturing, the culture was serially diluted with sterile phosphate buffer solution (PBS; pH 7.4), from which 0.1mL suspension was spread on the MSM-CIP-agar plate (CIP concentration = 0.5 g/L). The plates were incubated at 30 ^0^C for 24 h until the emergence of visible colonies. Single colony purification was done twice in MSM-CIP agar plates before being routinely maintained in slants. The purified culture was subsequently used for all physiological experiments including growth assays. Bergey’s Manual of Systematic Bacteriology (29) was followed to compare the results of biochemical tests. Genomic DNA was prepared and whole genome sequencing was done using the IlluminaNovaseq 6000 platform (30)). The whole genome sequence was used to construct the phylogenetic tree after aligning sequences of closely related type strains via the Type Strain Genome Server (https://tygs.dsmz.de/) (31).

### Growth assays in different media

The growth studies were conducted in Luria Bertani (LB) medium, a modified mineral salt medium (without N-source, as SG01 demonstrated growth in MSM without supplementation of any inorganic or organic form of nitrogen) supplemented with 5.0 g/L glucose (MSM-glucose) or 2 g/L ciprofloxacin (MSM-CIP) (media composition given in Supplementary file 1). Before performing detailed growth studies in LB or MSM-glucose, standard procedure was used to calibrate OD_600_ to colony forming unit (CFU) counts, which are directly relatable to the cell concentration of the culture, i.e. viable cell counts per mL. Following the protocol, 10 mL of overnight grown cells of a pre-culture in LB was centrifuged, pellet was re-suspended in 10 mL sterile PBS and serially diluted. From 10^−3^ diluted bacterial suspensions, cells were inoculated (1 %) into fresh sterile LB for enumerating growth using the standard plate technique. For conducting a growth assay in MSM-glucose, 10 mL of log phase grown cells (8 h grown culture) of a pre-culture in LB was centrifuged, and the pellet was re-suspended in 10 mL sterile PBS and diluted. From 10^−1^ diluted bacterial suspension, cells were inoculated (1 %) into fresh sterile MSM-glucose for enumerating growth using the standard plate technique. Similarly, for growth studies in MSM-CIP, 10 mL of log phase grown cells (7 h grown culture) of a pre-culture in MSM-glucose was centrifuged, and the pellet was re-suspended in 10 mL sterile PBS and diluted. From 10^−1^ diluted bacterial suspension, cells were inoculated (1 %) into fresh sterile MSM-CIP for enumerating growth using the standard plate technique.

### UV-vis Spectroscopic quantification of CIP

Spectrometric determination of residual CIP in the culture medium was done by using a Lambda 25 UV/VIS spectrometer. Absorption maxima (λ_max_) were determined before generating a standard curve using varying concentrations of CIP (10–90 μg/mL). For quantifying the concentration of CIP in the culture medium following inoculation of bacterium, the growing culture medium was withdrawn at specific intervals and centrifuged at 10,000 rpm to pellet down cells for collection of cell-free supernatant. The optical density of each cell-free supernatant sample (derived from samples at different time intervals) at λ_max_ was obtained to ascertain the CIP concentration from the standard curve (14).

### Residual Antibacterial activity of the degraded products

The residual antibacterial potential of intermediates formed in the course of degradation of CIP by SG01 was evaluated for inhibitory activity of culture medium minus cells (cell-free supernatant/ culture supernatant) against CIP-sensitive *Escherichia coli* K12 on Luria agar by disc diffusion method (32).

### Studying plasticity in cell dimensions of SG01 when grown in different media

SG01 was grown in four different media, LB, MSM-glucose, and MSM-CIP up to the mid-log phase. Cell harvesting time was ascertained from individual growth curves in a specific medium. Immediately before passing the log phase cells through a 0.22μm bacterial filter, unfiltered culture was serially diluted plated, and incubated overnight to determine the cell count. Side-by-side, the filtrate was also serially diluted and plated, and incubated overnight to determine the cell count (CFU/mL). Light microscopy of the log-phase cells grown in four different media was done following Gram staining (33) and negative staining using nigrosin (34). Scanning electron microscopy (SEM) was done to confirm a reduction in cell size when grown in MSM-CIP.

### Transcriptome study

Prior to transcriptome analyses, genomic DNA was isolated and sequenced as per the protocol described earlier (35). For transcriptome, the strain SG01 was grown in MSM-glucose (control) and MSM-CIP (test) media at 30 ^0^C under gyrotary shaking (100 r.p.m.). Cells were harvested from the mid-log phase by centrifugation at 4000 r.p.m. Spent media was discarded, and pelleted cells were washed twice with sterile PBS and immediately quenched with liquid nitrogen and stored at −80^0^C. Total RNA was isolated from both control and test bacterial samples, and after ascertaining qualities and quantities of the isolated RNA samples, RNA-seq sequencing libraries were prepared from the purified RNA samples using IlluminaTrueSeq mRNA sample Prep kit (Illumina, U.S.) and MICROBExpress kit (Invitrogen™, USA) as per the manufacturer’s protocol. Following sequencing, raw sequence reads were processed with Trimmomatic v0.39, which allowed for the removal of low-quality and adapter sequences from the raw data. The sequenced raw reads of the two samples, the control sample and test sample, were processed to obtain high-quality clean reads and were mapped on the reference genome of *Klebsiella* sp. SG01, using STAR (v 2.7.10a) with default parameters. Feature Counts (version 2.0.3) was used to count the number of reads mapped on each gene. Differential gene expression analysis was performed using the DEGSeqR package between control and test samples. Log2Fold change (log2 FC) values greater than zero were considered up-regulated whereas less than zero were down-regulated along the P-value threshold of 0.05 for significant results. Using KAAS (KEGG Automatic Annotation Server), functional annotations of every gene were performed against the curated KEGG genes database by the previously mentioned methods (36).

### Network analysis and Identification of the hub genes

The STRING database version 12.0 was used to construct a PPI network based on experimental results, automated text-mining, co-expression data, and other curated databases(37). The upregulated DEGs > 1 log 2FC and downregulated DEGs < −1 log 2FC were chosen to study the interaction. The PPI networks were functionally categorized using the CLueGO plugin of Cytoscape v3.8.0(38, 39). The hub genes and bottleneck genes were also deciphered by the Cytohubba plugin (36, 40)

## Statistical analysis

Statistical analysis was performed using GraphPad Prism (graphpad.com/quickcalcs/ttest/?format=SD). Test’ was performed to compare the mean of the two groups.

## Data availability

Raw sequence reads are available at SRA: PRJNA931810 under accessions SRR24804248 for the draft genome sequence of *Klebsiella* sp. SG01 and SRR29374586 for the whole transcriptome sequence.

## Supplementary Material

Supplemental material for this article is provided in Supplementary file 1, and two Excel files, Supplementary Excel file 1, and Supplementary Excel file 2 respectively.

## Acknowledgments

We would like to acknowledge the University of North Bengal, Raja Rammohanpur Campus, India-734010 for their support in conducting this study. We are indebted to the Department of Biotechnology, Government of India for funding a part of our work (BT/PR40383/BCE/8/1561/2020). S.G. is thankful to the Government of West Bengal (WBP211629117511) for providing financial aid.

S.G. participated in designing the experiments, performed the studies, analysed data, wrote and reviewed the manuscript; R.C. conceived the idea, designed the experiments, analysed data, and wrote and reviewed the manuscript.

## References

1. Khouja T, Mitsantisuk K, Tadrous M, Suda KJ. 2022. Global consumption of antimicrobials: impact of the WHO Global Action Plan on Antimicrobial Resistance and 2019 coronavirus pandemic (COVID-19). J Antimicrob Chemother 77:1491–1499.

2. Yang Q, Gao Y, Ke J, Show PL, Ge Y, Liu Y, Guo R, Chen J. 2021. Antibiotics: An overview on the environmental occurrence, toxicity, degradation, and removal methods. Bioengineered 12:7376.

3. Danner M-C, Robertson A, Behrends V, Reiss J. 2019. Antibiotic pollution in surface fresh waters: Occurrence and effects. Sci Total Environ 664:793–804.

4. Cherian JJ, Rahi M, Singh S, Reddy SE, Gupta YK, Katoch VM, Kumar V, Selvaraj S, Das P, Gangakhedkar RR, Dinda AK, Sarkar S, Vaghela PD, Bhargava B. 2021. India’s Road to Independence in Manufacturing Active Pharmaceutical Ingredients: Focus on Essential Medicines. 2. Economies 9:71.

5. Van Doorslaer X, Dewulf J, Van Langenhove H, Demeestere K. 2014. Fluoroquinolone antibiotics: an emerging class of environmental micropollutants. Sci Total Environ 500– 501:250–269.

6. O’Hagan D. 2008. Understanding organofluorine chemistry. An introduction to the C–F bond. Chem Soc Rev 37:308–319.

7. Zhao L, Dong YH, Wang H. 2010. Residues of veterinary antibiotics in manures from feedlot livestock in eight provinces of China. Sci Total Environ 408:1069–1075.

8. Rusu A, Hancu G, Uivarosi V. 2015. Fluoroquinolone pollution of food, water and soil, and bacterial resistance. Environ Chem Lett 13:21–36.

9. Gou N, Yuan S, Lan J, Gao C, Alshawabkeh AN, Gu AZ. 2014. A quantitative toxicogenomics assay reveals the evolution and nature of toxicity during the transformation of environmental pollutants. Environ Sci Technol 48:8855–8863.

10. Amorim CL, Moreira IS, Maia AS, Tiritan ME, Castro PML. 2014. Biodegradation of ofloxacin, norfloxacin, and ciprofloxacin as single and mixed substrates by Labrys portucalensis F11. Appl Microbiol Biotechnol 98:3181–3190.

11. Pan L-J, Li J, Li C-X, Tang X, Yu G-W, Wang Y. 2018. Study of ciprofloxacin biodegradation by a Thermus sp. isolated from pharmaceutical sludge. J Hazard Mater 343:59–67.

12. Nguyen LN, Nghiem LD, Oh S. 2018. Aerobic biotransformation of the antibiotic ciprofloxacin by Bradyrhizobium sp. isolated from activated sludge. Chemosphere 211:600–607.

13. Fang H, Oberoi AS, He Z, Khanal SK, Lu H. 2021. Ciprofloxacin-degrading Paraclostridium sp. isolated from sulfate-reducing bacteria-enriched sludge: Optimization and mechanism. Water Research 191:116808.

14. Ali Q, Zainab R, Badshah M, Sarwar W, Khan S, Mustafa G, Ibrahim T, Ahmed S. 2024. Prospecting the biodegradation of ciprofloxacin by Stutzerimonas stutzeri R2 and Exiguobacterium indicum strain R4 isolated from pharmaceutical wastewater. H2Open Journal 7:149–162.

15. Kajander EO, Ciftçioglu N. 1998. Nanobacteria: an alternative mechanism for pathogenic intra- and extracellular calcification and stone formation. Proc Natl Acad Sci U S A 95:8274–8279.

16. Abrol N, Panda A, Kekre NS, Devasia A. 2015. Nanobacteria in the pathogenesis of urolithiasis: Myth or reality? Indian J Urol 31:3–7.

17. Seymour CO, Palmer M, Becraft ED, Stepanauskas R, Friel AD, Schulz F, Woyke T, Eloe-Fadrosh E, Lai D, Jiao J-Y, Hua Z-S, Liu L, Lian Z-H, Li W-J, Chuvochina M, Finley BK, Koch BJ, Schwartz E, Dijkstra P, Moser DP, Hungate BA, Hedlund BP. 2023. Hyperactive nanobacteria with host-dependent traits pervade Omnitrophota. Nat Microbiol 8:727–744.

18. Raoult D, Drancourt M, Azza S, Nappez C, Guieu R, Rolain J-M, Fourquet P, Campagna B, La Scola B, Mege J-L, Mansuelle P, Lechevalier E, Berland Y, Gorvel J-P, Renesto P. 2008. Nanobacteria are mineralo fetuin complexes. PLoS Pathog 4:e41.

19. Fasnacht M, Polacek N. 2021. Oxidative Stress in Bacteria and the Central Dogma of Molecular Biology. Front Mol Biosci 8.

20. Szaleniec M, Wojtkiewicz AM, Bernhardt R, Borowski T, Donova M. 2018. Correction to: Bacterial steroid hydroxylases: enzyme classes, their functions and comparison of their catalytic mechanisms. Appl Microbiol Biotechnol 102:8173

21. Pervez MN, Ma S, Huang S, Naddeo V, Zhao Y. 2022. Photo-Fenton Degradation of Ciprofloxacin by Novel Graphene Quantum Dots/α-FeOOH Nanocomposites for the Production of Safe Drinking Water from Surface Water. 14. Water 14:2260.

22. Goo E, An JH, Kang Y, Hwang I. 2015. Control of bacterial metabolism by quorum sensing. Trends Microbiol 23:567–576.

23. Monahan LG, Turnbull L, Osvath SR, Birch D, Charles IG, Whitchurch CB. 2014. Rapid conversion of Pseudomonas aeruginosa to a spherical cell morphotype facilitates tolerance to carbapenems and penicillins but increases susceptibility to antimicrobial peptides. Antimicrob Agents Chemother 58:1956–1962.

24. Ojkic N, Banerjee S. Bacterial cell shape control by nutrient-dependent synthesis of cell division inhibitors - PubMed. https://pubmed.ncbi.nlm.nih.gov/33838134/. Retrieved 13 November 2024.

25. Cheng-Guang H, Gualerzi CO. 2021. The Ribosome as a Switchboard for Bacterial Stress Response. Front Microbiol 11.

26. Shrivastava M, Kaur M, Luke LM, Kakkar RA, Roy D, Singla S, Sharma V, Sharma G, Chaba R. 2024. An envelope stress response governs long-chain fatty acid metabolism via a small RNA to maintain redox homeostasis in Escherichia coli 10.1101/2024.10.18.618624.

27. Erhardt H, Steimle S, Muders V, Pohl T, Walter J, Friedrich T. 2012. Disruption of individual nuo-genes leads to the formation of partially assembled NADH:ubiquinone oxidoreductase (complex I) in Escherichia coli. Biochim Biophys Acta 1817:863–871.

28. Guest RL, Wang J, Wong JL, Raivio TL. 2017. A Bacterial Stress Response Regulates Respiratory Protein Complexes To Control Envelope Stress Adaptation. J Bacteriol 199:e00153–17.

29. 2000. Bergey’s manual of determinative bacteriology9. ed., [Nachdr.]. Lippincott Williams & Wilkins, Philadelphia.

30. Modi A, Vai S, Caramelli D, Lari M. 2021. The Illumina Sequencing Protocol and the NovaSeq 6000 System. Methods Mol Biol 2242:15–42.

31. Meier-Kolthoff JP, Göker M. 2019. TYGS is an automated high-throughput platform for state-of-the-art genome-based taxonomy. Nat Commun 10:2182.

32. Singh SK, Khajuria R, Kaur L. 2017. Biodegradation of ciprofloxacin by white rot fungus Pleurotus ostreatus. 3 Biotech 7:69.

33. Coico R. 2001. Gram staining. Curr Protoc Immunol Appendix 3:A.3O.1–A.3O.2.

34. Moyes RB, Reynolds J, Breakwell DP. 2009. Preliminary staining of bacteria: negative stain. Curr Protoc Microbiol Appendix 3:Appendix 3F.

35. Sen S, Saha T, Bhattacharya S, Nidhi null, Mondal N, Ghosh W, Chakraborty R. 2020. Draft Genome Sequences of Two Boron-Tolerant, Arsenic-Resistant, Gram-Positive Bacterial Strains, Lysinibacillus sp. OL1 and Enterococcus sp. OL5, Isolated from Boron-Fortified Cauliflower-Growing Field Soils of Northern West Bengal, India. Microbiol Resour Announc 9:e01438–19.

36. Sen S, Ganguli S, Chakraborty R. 2023. What transcriptomics and proteomics can tell us about a high borate perturbed boron tolerant Bacilli strain. Mol Omics 19:370–382.

37. Szklarczyk D, Kirsch R, Koutrouli M, Nastou K, Mehryary F, Hachilif R, Gable AL, Fang T, Doncheva NT, Pyysalo S, Bork P, Jensen LJ, von Mering C. 2023. The STRING database in 2023: protein-protein association networks and functional enrichment analyses for any sequenced genome of interest. Nucleic Acids Res 51:D638–D646.

38. Shannon P, Markiel A, Ozier O, Baliga NS, Wang JT, Ramage D, Amin N, Schwikowski B, Ideker T. 2003. Cytoscape: a software environment for integrated models of biomolecular interaction networks. Genome Res 13:2498–2504.

39. Bindea G, Mlecnik B, Hackl H, Charoentong P, Tosolini M, Kirilovsky A, Fridman W-H, Pagès F, Trajanoski Z, Galon J. 2009. ClueGO: a Cytoscape plug-in to decipher functionally grouped gene ontology and pathway annotation networks. Bioinformatics 25:1091–1093.

40. Chin C-H, Chen S-H, Wu H-H, Ho C-W, Ko M-T, Lin C-Y. 2014. cytoHubba: identifying hub objects and sub-networks from complex interactome. BMC Syst Biol 8 Suppl 4:S11.

